# Sub-clinical exposure to *Streptococcus pyogenes* drives the development of immunity

**DOI:** 10.1101/2025.07.02.662868

**Authors:** Ailin Lepletier, Despena Vedis, Vicky Ozberk, Merrina Anugraham, Darrell Bassette, Ainslie Callcutt, Hannah R. Frost, Kristy I. Azzopardi, Andrew C. Steer, Daniel Kolarich, Joshua Osowicki, Michael F. Good, Manisha Pandey

## Abstract

Age-related decline in *Streptococcus pyogenes* infection rates suggests that immunity develops progressively through repeated exposure during early life. However, the intensity or duration of exposure required is unknown, as to why some individuals appear to develop immunity, despite having few or no previously detected infections. Here, drawing on samples from a human challenge model of pharyngeal *S. pyogenes* infection, we investigate whether symptomatic disease is required for induction of humoral and cellular immunity. Challenge with M75 *S. pyogenes* induced M75-specific serum IgG and IgA antibodies and memory B cell in both symptomatic and asymptomatic participants, with responses persisting for at least 6 months. Purified IgG from asymptomatic participants exhibited significantly enhanced binding to M75 *S. pyogenes* and were bactericidal when transferred into a murine model of pharyngeal infection. M75-specific IgG from these participants had an altered Fc glycosylation signature, indicative of enhanced effector function and ability to limit inflammation. However, *S. pyogenes* challenge had no impact on cellular or humoral immune responses to a conserved cryptic epitope, p*17. These findings show that asymptomatic (or sub-clinical) exposure to M75 *S. pyogenes* generates functional immune responses and contributes to the streptococcal immunity that emerges by adulthood.

## Introduction

*Streptococcus pyogenes* (Group A *Streptococcus*, Strep A) is a leading human-restricted pathogen with a persistent and profound global disease burden. It causes broad clinical spectrum of illness, spanning superficial through to severe infections and post-infectious syndromes including rheumatic heart disease ^1,2^. Every year, up to a billion people are affected and more than 500,000 deaths are attributable to *S. pyogenes* ^3^. ‘Strep throat’ or *S. pyogenes* pharyngitis is a ubiquitous childhood illness, with incidence peaking during the primary school years ^4,5^. Lower incidence in adults has generally been ascribed to immunity accumulated through repeated exposure in childhood to different *S. pyogenes* strains ^6^. Although the precise mechanisms of immunity against *S. pyogenes* remain uncertain, several humoral and cellular responses have been strongly implicated in protection against colonization or disease ^7^. These responses include antibodies that bind to bacteria, enabling neutrophils and monocytes at the site of infection to recognize and eliminate them through opsonophagocytosis ^7^. In individuals resistant to pharyngeal acquisition, such antibodies can also prevent early bacterial colonization by blocking adherence and promoting rapid clearance^8^.

Allelic polymorphism of the *S. pyogenes* M-protein is a major impediment to the development of immunity. To date, over 250 distinct serotypes have been identified based on its amino-terminal hypervariable region (HVR) ^9^. Longitudinal studies indicate that naturally acquired antibodies against the M-protein can provide homologous protection but offer limited cross-protection against disease caused by heterologous strains ^10,11^. While the M cluster concept explains some degree of cross-protective immunity ^12^, it remains unclear how most children develop protective immunity by their second decade of life ^12^, considering that only around 15% experience symptomatic *S. pyogenes* pharyngitis in any given year ^1^. A potential explanation is that repeated exposures boost immunity to antigens that are broadly conserved across strains, complementing narrower responses to type-specific antigens such as the M-protein HVR. We previously showed that antibodies to the conserved C-repeat region can kill the *S. pyogenes* irrespective of its M-protein serotype ^13–15^. However, C-repeat-specific immunity is unlikely to account for naturally acquired immunity because this region is poorly immunogenic (‘cryptic’) during both experimental and natural infections. Antibodies to this region do not develop in children from streptococcal-endemic areas until well into their second decade of life ^16^ nor do they arise following repeated exposures of mice to *S. pyogenes*^17^.

Given the high prevalence of asymptomatic pharyngeal *S. pyogenes* colonization or ‘carriage’ in 10–20% of school-age children ^18^ and that many different serotypes can be in circulation in a community at any time ^19^, a reasonable hypothesis is that repeated asymptomatic (or sub-clinical) exposures shape protective immune responses to *S. pyogenes*. Here, we explore this hypothesis in a well-characterised cohort of participants from the CHIVAS-M75 *S. pyogenes* human pharyngeal challenge study ^20–22^. While acute symptomatic pharyngitis following direct tonsillar inoculation was the most common outcome, a subset remained asymptomatic despite lacking significant pre-existing serotype-specific antibodies to the M75 strain, enabling us to investigate the immunological outcomes of sub-clinical infections. At 6-months post-challenge, increased levels of memory B cells and antibodies targeting the M75 hypervariable region (HVR) —but not a conserved cryptic M-protein epitope (p*17) —were observed in participants both with and without pharyngitis. Using several approaches to assess antibody function, including bacterial binding, IgG Fc glycosylation traits, and passive antibody transfer into a murine model of pharyngeal infection, we show that sub-clinical exposures can elicit immune responses that contribute to long-term protection against *S. pyogenes*.

## Results

### Murine antibodies against the M75 hypervariable region bind specifically to M75 *S. pyogenes*

Our initial studies aimed to characterize humoral immune responses elicited by the M-protein peptides, p*17 and M75-HVR (Fig 1A). Although p*17 is poorly recognized following infection of mice, it is known that vaccine-induced antibodies can protect against infection ^23–26^. BALB/c mice were immunized with either p*17-diphtheria toxoid (DT) or M75-HVR-DT, each adjuvanted with aluminum hydroxide (Alum), following a 3-dose schedule as in a Phase 1 clinical trial ^27^. Serum from both groups were collected two weeks after the final vaccine dose and analyzed by indirect ELISA and flow cytometry-based binding assays to assess IgG antibody responses against p*17 and M75-HVR peptides, as well as to live *S. pyogenes* bacteria (Fig 1B), respectively. Serum from M75-DT immunized mice specifically recognized the M75 peptide (Fig. 1C), whereas serum from p*17-DT-immunized mice exclusively recognized p*17 (Fig. 1D). Using flow cytometry, we demonstrated that serum from M75-DT–immunized mice specifically bound to M75 *S. pyogenes*, with no detectable reactivity to an M1 strain (Fig. 1E). In contrast, serum from p*17-DT–immunized mice recognized both M75 and M1 strains (Fig. 1F), consistent with p*17 being a conserved epitope across *S. pyogenes* strains.

**Figure 1.**
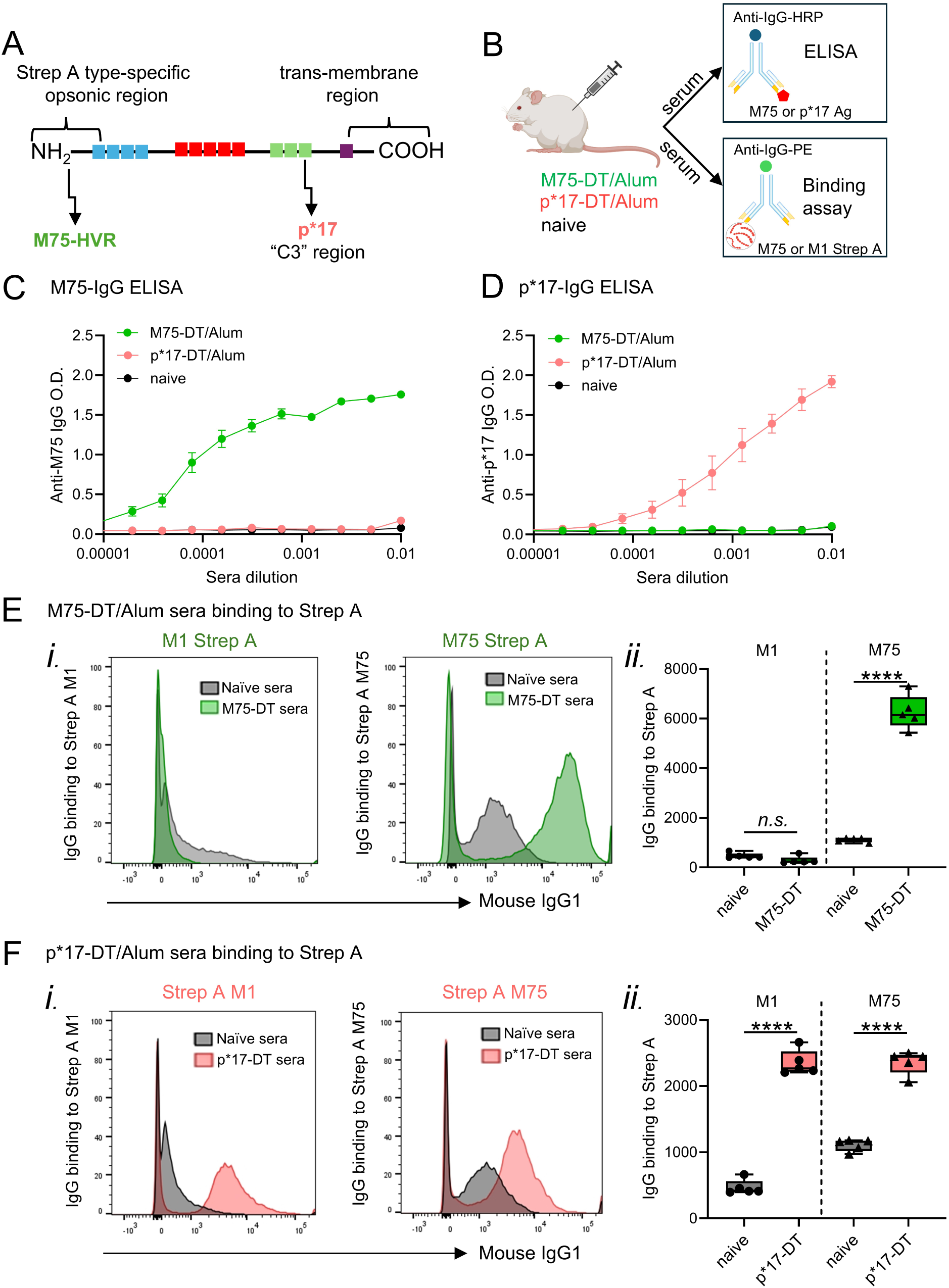
Domain localization, antibody responses, and *S. pyogenes* binding following immunization with M75-HVR and p*17 epitope-based vaccines. (A) Schematic representation of M-protein domains indicating the N-terminal position of M75 epitope, which is derived from the HVR region of the M-protein (M75-HVR), and the highly conserved p*17 epitope, derived from the C-terminal (C3) position. (B) Sera collected two weeks after the final immunization of BALB/c mice with p*17-DT or M75-DT adjuvanted with Alum (n=5/group) were assessed using direct ELISA and *S. pyogenes* binding assays. (C and D) ELISA was used to detect (C) anti-M75 and (D) anti-p*17-specific IgG at O.D. 450 nm. Absorbance values are shown for sera collected from mice immunized with M75-DT/Alum, p*17-DT/Alum or naïve controls at multiple dilutions. (E and F) Flow cytometry-based assay analyzing the binding of sera collected from mice immunized with (E) M75-DT/Alum or (F) p*17-DT/Alum to M1 and M75 live *S. pyogenes*. Serum obtained from naïve mice was used as controls. i. Representative histograms illustrating *S. pyogenes* binding by sera from one mouse per group. ii. Box plots depicting IgG1 antibody binding to *S. pyogenes*, quantified as mean fluorescence intensity (MFI). Unpaired t-tests were used for statistical analysis. n.s. non-significant; ****P<0.0001

### Human pharyngeal challenge with the M75 *S. pyogenes* elicits memory responses directed against the M-protein hypervariable region

To assess the long-term human immune responses to infection, we examined memory B cell populations and serum antibodies from CHIVAS-M75 participants ^20–22^. Antigen-specific responses to p*17 and M75-HVR were assessed for 13 participants who developed acute symptomatic pharyngitis following pharyngeal challenge with M75 *S. pyogenes*. These were compared to responses from 5 participants who remained asymptomatic (non-pharyngitis) and had no detectable M75 *S. pyogenes* by culture or qPCR from throat swabs collected twice daily from 24 hours after challenge until discharge. Systemic immune responses were longitudinally analyzed at pre-challenge, 1 month, and 6 months post-challenge.

To evaluate immunity against *S. pyogenes* and explore the dynamics of antigen-specific memory B cells and antibody responses post-challenge, we generated fluorescent tetramers and analyzed responses to the p*17 and the type-specific HVR region of the M75 protein. To determine whether a single M75 *S. pyogenes* infection induced antibodies to the conserved region of the M-protein, we specifically assessed responses to p*17 (Fig. 2A and B). Flow cytometry-based tetramer analysis was used to characterize p*17-specific memory B cells (CD27^+^CD21^+^) within both the IgA and IgG subclasses (Supp. Fig. 1A). The frequency of p*17-specific IgG^+^ memory B cells remained low and unchanged following pharyngeal challenge with M75 *S. pyogenes* out to 6 months in both symptomatic and asymptomatic participants (Fig. 2A). Likewise, p*17-specific IgA^+^ cells in the blood were unaffected by the challenge, with no detectable link to disease progression (Fig. 2B). Antibody response analyses in the same participants similarly revealed that IgG antibody levels targeting p*17 remained low and unchanged across all timepoints in all participants following pharyngeal challenge (Fig. 2C). Similarly, serum IgA responses to the p*17 antigen were minimal (O.D. < 0.2 at a dilution of 1:20), showing no significant changes post-challenge and no association with disease progression (Fig. 2D).

**Figure 2.**
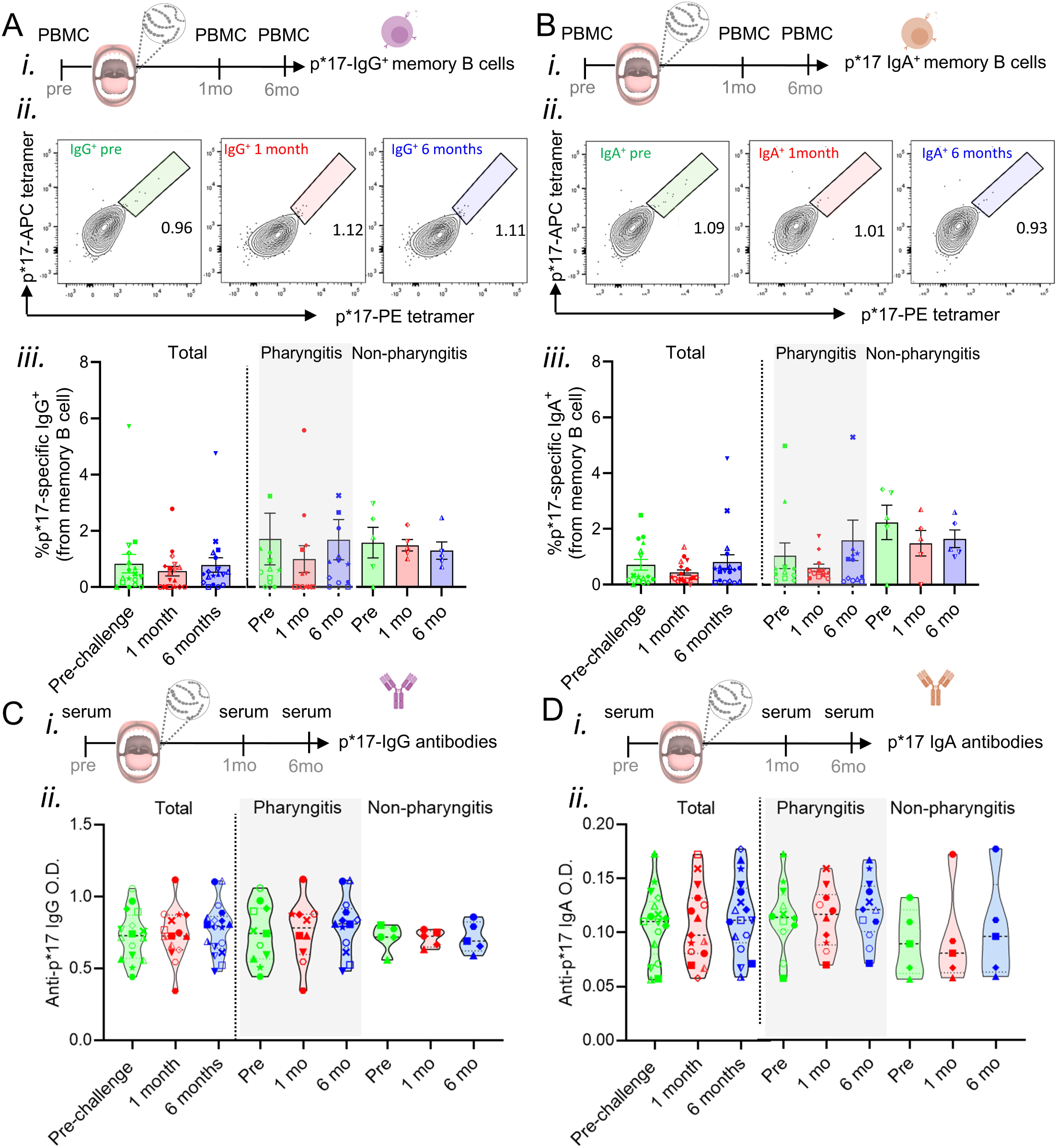
Memory responses to p*17 following human infection with *S. pyogenes* M75. (A, B). Flow cytometry analysis of p*17-specific (A) IgG+ and (B) IgA+ memory B cells in the peripheral blood of participants in a controlled human infection study. i. Schematic overview of study timeline and collection of peripheral blood mononuclear cells (PBMC) for analysis of B cells in the peripheral blood. ii. Flow cytometry gating strategy for analysis of antigen-specific memory B cells in the peripheral blood. iii. The percentage of p*17-specific memory B cells is shown in the bar graphs at pre-challenge (day -1), 1-month post-challenge, and 6 months post-challenge. (C, D) ELISA was used to detect anti-p*17-specific (C) IgG and (D) IgA antibodies in PBMC-matched serum samples. i. Schematic overview of study timeline and collection of serum for analysis of antibodies. ii. Violin plots show O.D. at 450 nm absorbance values for sera collected at pre-challenge (day -1) and 6 months post-challenge, using a 1:20 dilution. Data are stratified by participants with and without pharyngitis. Each symbol represents an individual participant. Statistical analysis was performed using one-way ANOVA followed by Bonferroni’s multiple comparisons test. No significant differences were observed between the groups.

In those who developed symptomatic disease post-challenge, the frequency of M75-specific IgG^+^ memory B cells increased significantly at 6 months (Fig. 3A). M75-specific IgA^+^ memory B cells showed an early increase, observed at 1 month, which also endured for at least 6 months post-challenge (Fig. 3B). In asymptomatic participants there was a trend toward an increase at 6 months for both M75-specific IgG^+^ (2.09 fold-increase, P>0.05, Fig. 3A) and IgA^+^ (1.69 fold-increase, P>0.05, Fig. 3B) memory B cells.

**Figure 3.**
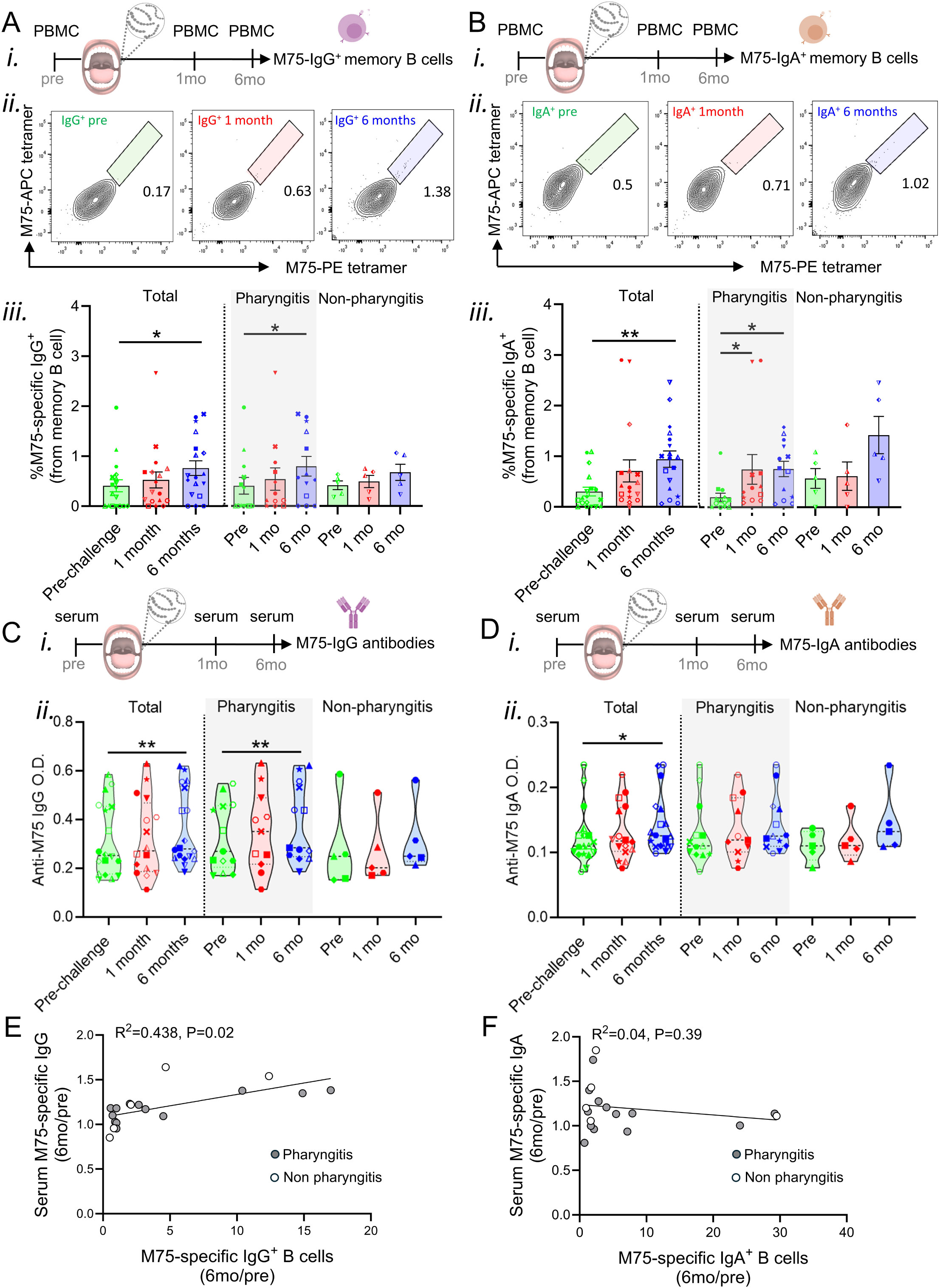
Memory responses to M75-HVR following human infection with *S. pyogenes* M75. (A, B) Flow cytometry analysis of M75-specific (A) IgG+ and (B) IgA+ memory B cells in the peripheral blood of participants in a controlled human infection study. i. Schematic overview of study timeline and collection of PBMC for analysis of B cells. ii. Flow cytometry gating strategy for analysis of antigen-specific memory B cells in the peripheral blood. iii The percentage of M75-specific memory B cells is shown in the bar graphs at pre-challenge (day -1), 1-month post-challenge, and 6 months post-challenge. (C, D) ELISA was used to detect anti-M75-specific (C) IgG and (D) IgA antibodies in PBMC-matched serum samples. i. Schematic overview of study timeline and collection of serum for analysis of antibodies. ii. Violin plots show at O.D. 450 nm absorbance values for sera collected at pre-challenge (day -1) and 6 months post-challenge, using a 1:20 dilution. Data are stratified by participants with and without pharyngitis. Each symbol represents an individual participant. Statistical analysis was performed using one-way ANOVA followed by Bonferroni’s multiple comparisons test. *P<0.05, **P<0.01. (E, F) Correlation between the fold change in M75-specific (E) IgG+ and (F) IgA+ memory B cells and the fold change in serum M75-specific antibodies at 6 months post- vs pre-challenge. Full circles = pharyngitis, empty circle non-pharyngitis participants. Correlation coefficient and P value were determined using Spearman’s method with a linear regression line included to indicate linear trends.

Similarly, antibody analysis showed a rise in M75-specific IgG levels in participants with pharyngitis but was detected only at 6 months post-challenge (Fig. 3C). This increase was driven specifically by the M75-specific IgG3 subtype (Supp. Fig. 1B) with no significant changes in IgG1, IgG2, or IgG4 levels (Supp. Fig. 1C-E). M75-specific serum IgA antibodies also showed a significant increase detected at 6 months post-challenge (Fig. 3D). Although a significant antibody increase was observed only in symptomatic participants, the average increase in M75-specific antibody levels was similar in those with and without pharyngitis (Supp. Fig. 1F) (Total IgG: mean values of 1.176 for pharyngitis and 1.242 for non-pharyngitis; IgG3: mean values of 1.431 for pharyngitis and 1.435 for non-pharyngitis; IgA:1.147 for pharyngitis and 1.330 for non-pharyngitis). The increase in IgG^+^ memory B cells binding the M75 antigen correlated with increased M75-specific serum IgG at 6 months relative to pre-challenge (R^2^=0.438, P=0.02, Fig. 3E). Interestingly, the increase in IgA^+^ memory B cells binding the M75 *S. pyogenes* antigen did not correlate with increased M75-specific serum IgA (R^2^=0.04, P=0.39, Fig. 3F). There was no significant difference in pre-challenge M75-specific memory B cells and antibodies between participants who did and did not develop pharyngitis following challenge (Fig. 3C and D).

Exposure to M75 *S. pyogenes* induces long-term, strain-specific systemic immunity, as evidenced by sustained serum Ig levels and memory B cell responses involving both IgG and IgA. While occurring in both groups of participants, the immune responses were more pronounced in those who developed clinical disease.

### IgG Fc glycosylation is associated with clinical manifestations of *S. pyogenes* infections

Post-translational modifications, particularly glycosylation of the constant (Fc) IgG domain, are well-known features of all IgGs and play a critical role in modulating antibody effector functions, including complement activation and Fc receptor engagement ^28,29^.

We initially assessed whether pre-challenge IgG Fc N-glycosylation traits could be a distinguishing factor between participants with and without pharyngitis. We affinity-purified total (non-specific) and M75-specific IgG from asymptomatic and symptomatic participants prior to challenge. M75-specific IgG levels pre-challenge were very low in all participants, but affinity purification enabled enrichment of these antibodies (Fig. 3C and 4A). Mass spectrometry-based characterisation revealed that pre-challenge IgG1 antibodies were the major IgG subclass targeting the M75-epitope across all participants. Participants who remained asymptomatic following challenge displayed a distinct baseline Fc glycosylation profile for global (entire IgG pool present in plasma) IgG1 compared to participants who became symptomatic (1.5-fold and 2.16 fold-increase in sialylated and afucosylated N-glycans, P<0.01 and P>0.05, respectively, Fig. 4B, *i*), whereas no significant differences were detected in overall glycoform distribution (Fig. 4C, *i*). Profound changes were observed in the glycosylation profile of M75-specific IgG1 antibodies from asymptomatic participants, marked by a 30-fold decrease in afucosylation (P<0.05) and a 4.9-fold increase in bisecting GlcNAc moieties (P<0.01, Fig. 4D, *i*), which is largely derived from increased levels of G1FB and G2FB N-glycans (3.1-fold and 13.9-fold increase, P<0.05 and P<0.01, respectively, Fig. 4E, *i*), compared to M75-specific IgG1 from symptomatic participants. In comparison to global IgG1, M75-specific IgG1 in participants without pharyngitis displayed significantly lower levels of afucosylated glycans (mean of 0.12 versus 2.13; P<0.01), along with a significantly higher proportion of bisected glycans (mean of 5.29 versus 1.36; P< 0.01, Figure 4B, *i* and D, *i*, respectively).

**Figure 4.**
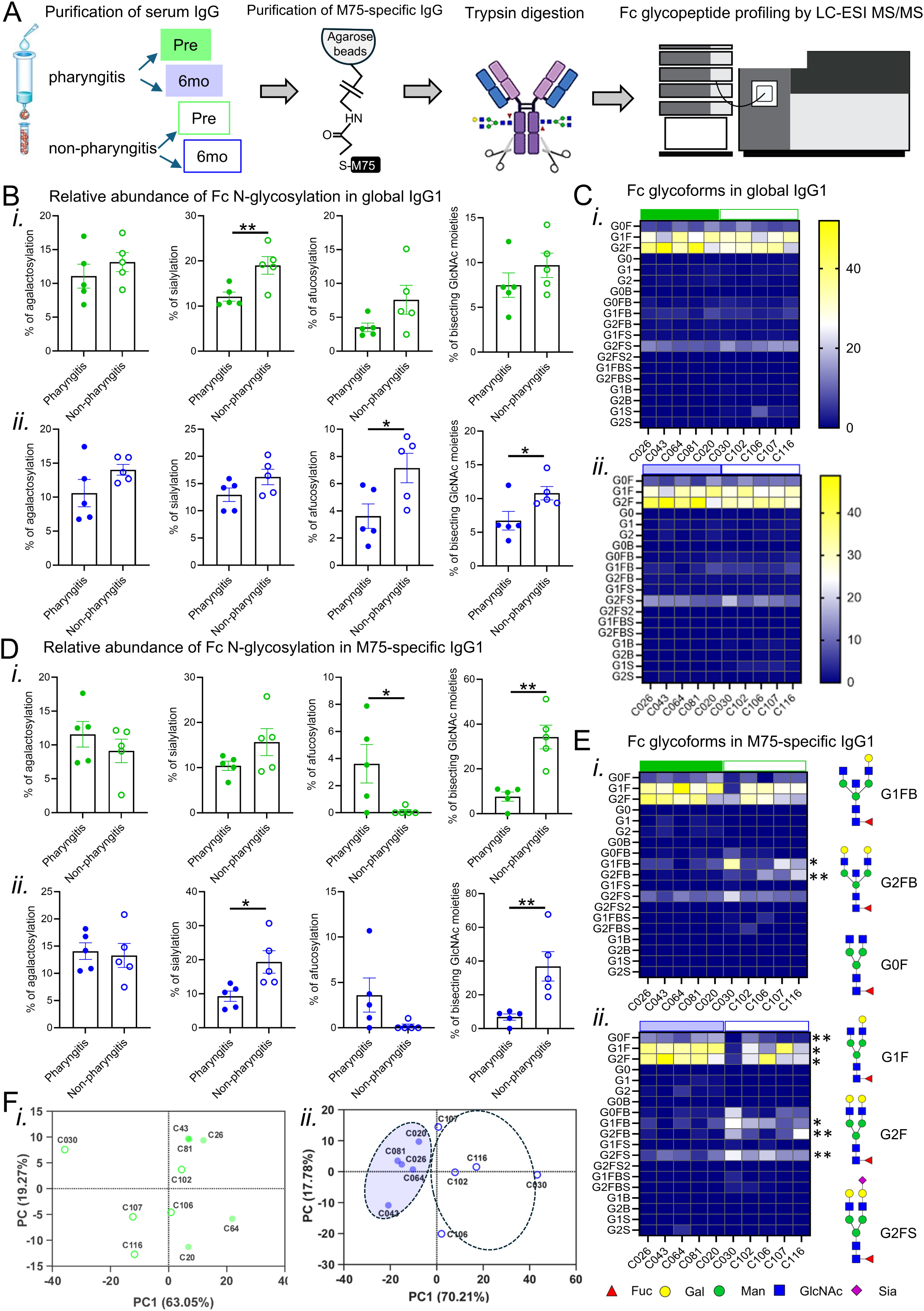
Comparative profiling of IgG Fc N-glycosylation in global and M75-specific serum IgG1 from participants with and without pharyngitis at pre-challenge and 6 months post-challenge. (A) Schematic overview of M75-specific IgG purification, digestion and Fc glycan profiling by liquid chromatography tandem mass spectrometry (LC-MS/MS). (B) Bar graphs depict the relative abundance of glycans in total IgG1 antibodies at *i.* pre-challenge and *ii.* 6 months post-challenge. (C) Heatmaps illustrate the fractional distribution of individual Fc glycoforms in global IgG1 antibodies at *i.* pre-challenge and *ii.* 6 months post-challenge. (D) Bar graphs depict the relative abundance of total glycans in M75-specific antibodies at *i.* pre-challenge and *ii.* 6 months post-challenge. (E) Heatmaps illustrate the fractional distribution of individual Fc glycoforms in M75-specific IgG1 antibodies at *i.* pre-challenge and *ii.* 6 months post-challenge. Schematic representation of the Fc glycoforms attached to IgG1 differentially distributed across participants with and without pharyngitis. Data represent individual unpaired samples from participants with (full symbols) and without (empty symbols) pharyngitis collected at pre-challenge (green) and 6 months-post challenge (blue) (n = 5/group). Unpaired t-test between participants with and without pharyngitis was used for statistical analysis *P<0.05, **P<0.01. (F) Principal component analysis shows the relationship of Fc glycoforms data from 5 biological replicates across participants with and without pharyngitis collected at *i.* pre-challenge (green) and *ii.* 6 months-post challenge (blue).

IgG Fc N-glycosylation traits at 6 months post-challenge showed similar differences between participants with and without pharyngitis as those observed pre-challenge. Participants who remained asymptomatic following challenge exhibited a 1.31-fold increase in sialylated N-glycans, a 1.98-fold increase in afucosylated N-glycans, and a 1.65-fold increase in bisecting GlcNAc moieties (P > 0.05, P < 0.05 and P < 0.05, respectively; Fig. 4B, *ii*), whereas no significant differences were detected in overall glycoform distribution (Fig. 4C, *ii*). Profound changes were observed in the glycosylation profile of M75-specific IgG1 antibodies from asymptomatic participants, marked by a 17.3-fold decrease in afucosylation (P>0.05), a 5.4-fold increase in bisecting GlcNAc moieties (P<0.01), and a 1.3-fold increase in sialylation (P<0.05, Fig. 4D, *ii*). This is derived not only from increased levels of G1FB, G2FB and G2FS N-glycans (3.9-fold, 11.3-fold and 2.1-fold increase, P<0.05, P<0.01 and P<0.01, respectively), but also from decreased levels of G0F, G1F and G2F (2.5-fold, 1.8-fold and 1.6-fold decrease P<0.01, P<0.05 and P<0.05, respectively, Fig. 4E, *ii*) compared to M75-specific IgG1 from symptomatic participants. The evident differences at post-challenge were further supported by principal component (PC) analyses, which revealed that Fc glycoforms profiles obtained from symptomatic and asymptomatic participants formed two well-defined and distinct clusters (Fig. 4F, *ii*).

Compared to pre-challenge levels, no significant differences were detected in overall IgG Fc N-glycosylation traits or glycoform compositions of global and M75-specific IgG1 antibodies from participants with pharyngitis at 6 months post-*S. pyogenes* challenge (Supp. Fig. 2). No changes in IgG Fc N-glycosylation were observed in global IgG1 purified from asymptomatic participants at 6 months post-challenge compared to pre-challenge levels either (Figures 5A-C). However, at 6 months, M75-specific IgG1 showed a significant increase in agalactosylation (1.5-fold) and sialylation (1.3-fold, P<0.05, Figure 5D) followed by a specific increase in G2FS (Figure 5E) in comparison to pre-challenge.

**Figure 5.**
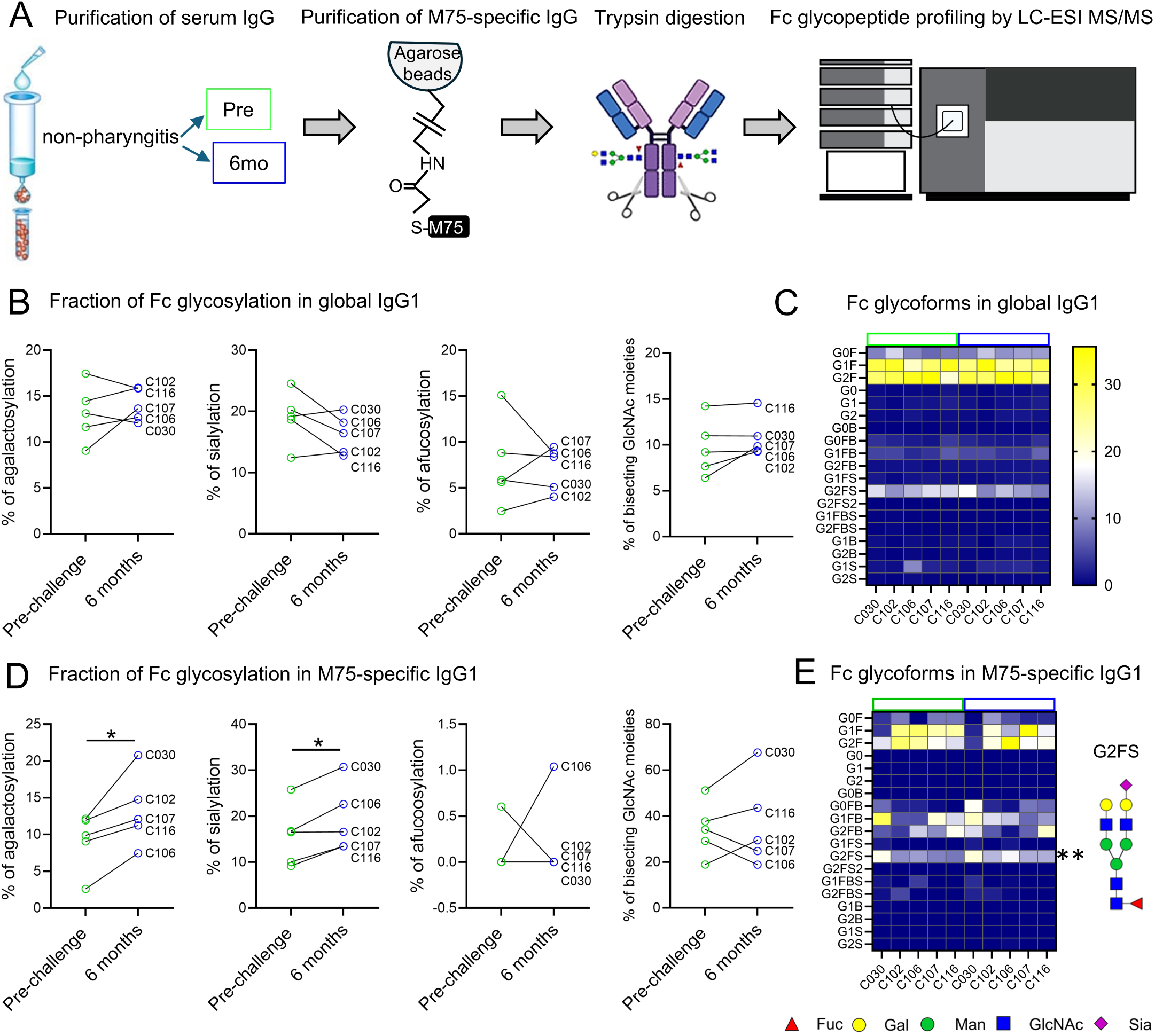
Fc N-glycosylation profiling of global and M75-specific serum IgG1 pre- and 6 months post-challenge in asymptomatic participants. (A) Schematic overview of M75-specific IgG purification, digestion and Fc glycan profiling by LC-MS/MS. (B) Line graphs depict the relative abundance of glycans in global IgG1 antibodies. (C) Heatmaps illustrate the fractional distribution of individual Fc glycoforms in global IgG1 antibodies. (D) Line graphs depict the relative abundance of glycans in M75-specific IgG1 antibodies in each participant. (E) Heatmaps illustrate the fractional distribution of individual Fc glycoforms in M75-specific IgG1 antibodies. Data represent paired samples collected at the time of challenge (green) and 6 months post-challenge (blue) from participants without pharyngitis (n = 5/group). Paired t-test between pre- and post-challenge was used for statistical analysis *P<0.05, **P<0.01.

These findings strongly indicate that the Fc-glycosylation traits of IgG1 antibodies directed against the M75 antigen are involved in modulating the susceptibility to symptomatic disease following M75 *S. pyogenes* challenge.

### Sub-clinical exposure to *S. pyogenes* induces long-term, functionally protective IgG immunity

To examine the functional properties of serum IgG and to increase the sensitivity of our analyses, we purified total serum IgG and used flow cytometry to assess the *S. pyogenes* binding at both pre-challenge and 6 months post-challenge in the participants with and without pharyngitis (Fig. 6A). We observed a 12.1-fold increase in the binding of purified IgG to the homologous M75 *S. pyogenes* at 6 months post-challenge in participants with symptomatic pharyngitis, relative to their pre-challenge levels (Fig. 6B). We also observed a significant increase in IgG binding (6.4-fold, P<0.05) in asymptomatic participants comparing 6-month post challenge relative to their pre-challenge levels (Fig. 6B). Similar to sera from mice immunized with M75-DT, IgG from CHIVAS trial participants at 6 months post-challenge (in both pharyngitis and non-pharyngitis sufferers) showed no binding to the heterogeneous M1 *S. pyogenes* (Figures 1E and 6C). These findings indicate that infection with *S. pyogenes* can elicit a significant rise in serotype-specific antibodies that persist for at least 6 months regardless of the clinical outcome.

**Figure 6.**
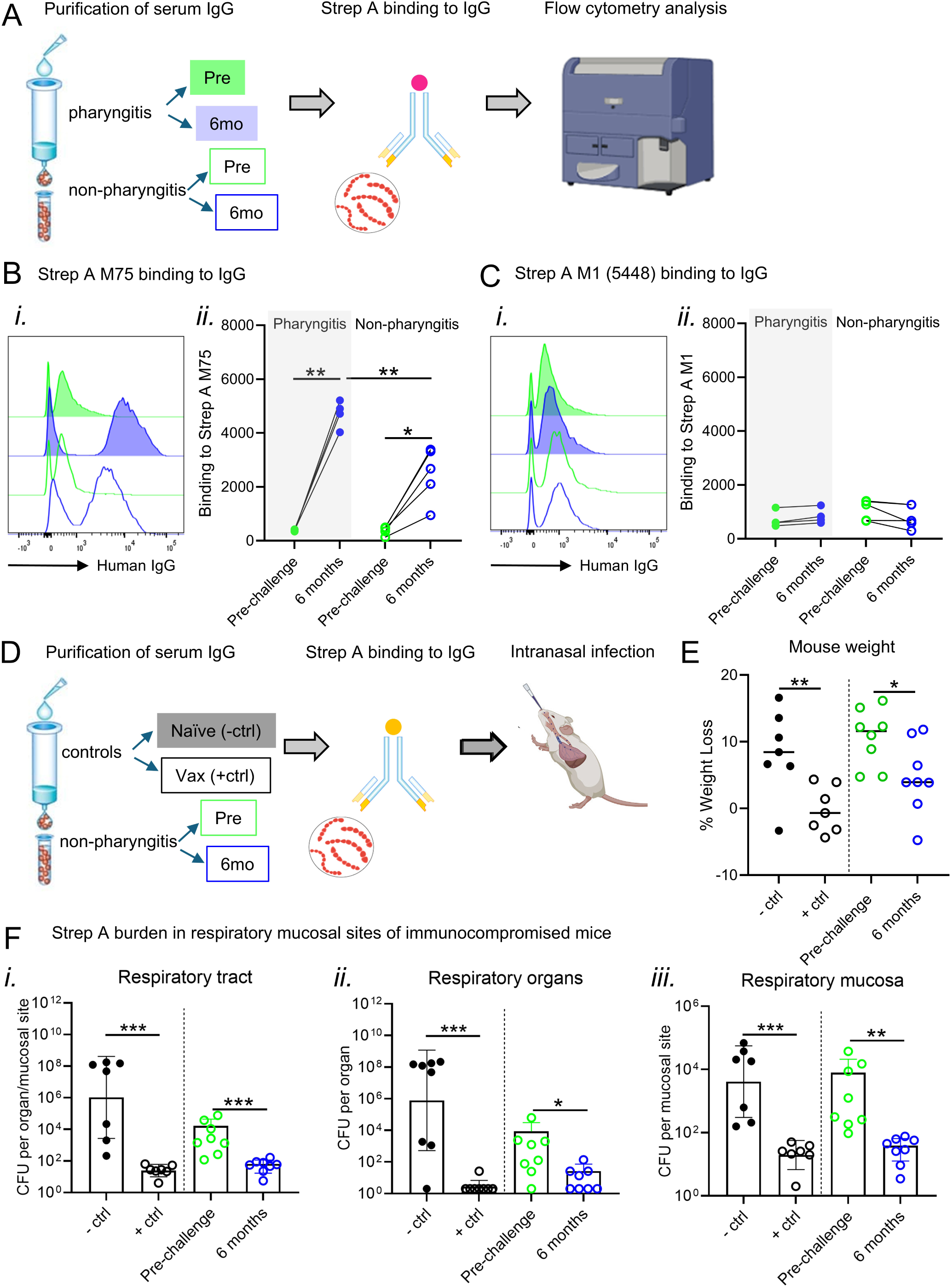
Functional analysis of purified serum IgG following *S. pyogenes* binding. (A-C) Flow cytometry-based binding assay shows the binding of IgG purified from sera to (B) M75 and (C) M1 *S. pyogenes*. Samples were collected at pre-infection (green) and 6 months post-infection (blue) from participants with (filled circles, n=5) and without (empty circles, n=5) pharyngitis. Each circle represents one participant. The level of binding of human IgG antibodies to *S. pyogenes* is shown as mean fluorescence intensity (MFI). One-way ANOVA followed by Bonferroni’s comparisons tests were performed in all statistical analyses. *P<0.05, **P<0.01. (D-F) SCID mice (n=7-8/group) were inoculated intranasally with M75 *S. pyogenes* incubated with pooled IgG serum from asymptomatic participants (n=5) at pre-challenge and 6 months post-challenge. IgG purified from rat sera immunized with p*17-K4S2-CRM/Alum or left as naïve was used as a control. (E) Graph shows the percentage of weight loss on day 2 following inoculation. (F) *S. pyogenes* colony-forming units (CFU) were enumerated in (i) total respiratory tract (total CFU obtained from throat swabs, nasal shedding, lungs and nasal-associated lymphoid tissue (NALT) throat swabs, (ii) in mucosal sites (throat swabs and nasal shedding) and (iii) in respiratory organs (CFU obtained lungs NALT and lungs), on day 2 post-challenge. An unpaired t-test was performed to compare weight loss and bacterial burden between mice treated with IgG purified from vaccinated versus naïve rats and from humans at 6 months post-challenge versus pre-challenge. n.s., non-significant; *P<0.05; **P<0.01; ***P<0.005.

The temporal changes observed in the IgG Fc N-glycosylation of asymptomatic participants prompted us to examine their functional properties more closely. We utilized a recently developed *in vivo* passive transfer model to test functional immunity. In this assay, *S. pyogenes* organisms were incubated with serum IgG antibodies (’pre-opsonized’) and then inoculated intranasally into immunocompromised mice. Clinical signs and bacterial burden in the respiratory tract were subsequently assessed to determine the protective capacity of the antibodies ^24^. Due to limited availability of serum, we purified IgG from the 5 asymptomatic participants at pre and 6-months post challenge, pooled the IgG at each time point, and tested it against M75 *S. pyogenes* in 7-8 mice for each group (Figure 6D). Further controls included IgG from rats vaccinated with p*17-K4S2-CRM ^26^ or IgG from naïve rats. We tested weight loss (as a general measure of disease in *S. pyogenes* -infected mice ^24^), and bacterial burden from the entire respiratory tract. Mice inoculated with M75 *S. pyogenes* that had been pre-incubated with IgG from vaccinated rats exhibited significantly less weight loss compared to mice administered *S. pyogenes* treated with IgG from naive rats (0% versus 8.4%, P<0.01) (Figure 6E). Across all mucosal sites and organs analyzed for bacterial burden, IgG from p*17-vaccinated rats resulted in significant reductions (P<0.01 – P<0.005) (Figure 6F).

We similarly observed that mice inoculated with M75 *S. pyogenes* pre-incubated with IgG from CHIVAS-M75 participants collected 6 months post-challenge exhibited significantly reduced weight loss compared to those administered with *S. pyogenes* incubated with pre-challenge IgG (4.64% versus 10.63%, P<0.05) (Figure 6E). This correlated with 273-fold reduction in total *S. pyogenes* colonization of the respiratory tract (P<0.005), which included both mucosal sites (nasal shedding and throat swabs) and respiratory organs (nasal-associated lymphoid tissue [NALT] and lungs). Specifically, we observed a 205-fold reduction in *S. pyogenes* colonization of the respiratory mucosa (P<0.01) and 314-fold reduction in colonization of respiratory tissues (P<0.05). There were no significant differences observed between IgG from naïve rats and the human pre-challenge IgG (Figure 6F).

Thus, serum IgG antibodies generated from sub-clinical *S. pyogenes* infection exhibit both strong strain-specific binding and robust functional activity.

## Discussion

This study demonstrates that functional immunity to *S. pyogenes* can develop following asymptomatic exposure. Pharyngeal human challenge with M75 *S. pyogenes* elicited class-switched B cells and antibodies (IgG and IgA) in the peripheral blood targeting the type-specific M-protein HVR region. While the immune responses elicited by asymptomatic challenge were lower than those observed in symptomatic participants, IgG from these participants demonstrated clear strain-specific binding and effectively mediated bacterial clearance in a passive transfer murine infection model, whereas pre-challenge IgG did not. Participants with higher frequency of M75-specific IgG memory B cells at 6 months after challenge also had higher levels of binding anti-M75 IgG, suggesting that memory B cells may help maintain or increase IgG antibody levels over time. These results support the hypothesis that immunity to *S. pyogenes* is gradually acquired throughout childhood by repeated asymptomatic (or sub-clinical) exposures to circulating strains and not only through symptomatic infections.

Recent longitudinal studies have also highlighted that asymptomatic *S. pyogenes* exposures may be immunologically significant (*i.e.* capable of triggering measurable immune responses), challenging the long-held assumption that only symptomatic infections contribute meaningfully to immunity. Studies in the USA by Hysmith et al. ^30^ and in The Gambia by Keeley et al. ^31^ each demonstrated antibody binding responses to M and non-M-protein antigens following asymptomatic episodes of *S. pyogenes* colonization. This observation is confirmed by other studies showing that a significant proportion of people with asymptomatic colonization of the upper respiratory tract develop functional anti-M-protein antibodies ^32,33^. Notably, a significant fraction of the children assessed in these studies only developed M-type specific serum IgG antibodies between 6 weeks and a year from detection of a positive throat culture ^30,32^, which aligns with our data showing a delayed immune response following pharyngeal challenge. In the 2-year study by Hysmith et al., new *S. pyogenes* acquisitions did not significantly boost homologous M-type-specific antibody responses ^30^. Therefore, it is possible that pre-screening and exclusion of participants with high serum type-specific M75 IgG in the CHIVAS-M75 study ^21,22^ may have contributed to the observed immunologically significant M75-specific responses following pharyngeal challenge.

Serum antibodies against the M-protein enhance the opsonisation and killing of *S. pyogenes* and are believed to contribute to immunity against homologous strains. We previously observed that a single intranasal infection with *S. pyogenes* resulted in long-lasting immunity against subsequent homologous challenge in a mouse model ^34^. Here we showed that human serum IgG antibodies, purified 6 months post-challenge, recognised the strain-specific HVR, bound specifically to live M75 *S. pyogenes*, and were bactericidal. The anti-M75 IgG response observed in the present study is predominantly mediated by the IgG1 and IgG3 subclasses, which are particularly effective in pathogen clearance, presenting a number of antigen-specific effector functions, including interaction of antigen-clustered IgG with the complement system, which activates phagocytosis and production of reactive oxygen species by innate immune cells ^35–37^. Despite its relatively low abundance in relation to IgG1, IgG3 has been increasingly recognized as a key player in the immune defence against various pathogens, including *S. pyogenes* ^38^ due to its superior affinity to Fc-gamma receptors ^39^. However, excessive IgG3 recognising M-protein antigens may be detrimental and elevated levels have been linked to the *S. pyogenes* post-infectious syndrome acute rheumatic fever^40,41^.

Although total and subclass-specific M75-IgG antibody levels prior to challenge did not distinguish between clinical outcomes following experimental challenge, significant differences in IgG Fc N-glycosylation traits both in global and M75-specific IgG1 were observed. Compared to those with symptomatic pharyngitis, asymptomatic participants had higher total IgG1 sialylation, suggesting that the anti-inflammatory properties of Fc sialylation ^42,43^ may also contribute to protection. Additionally, they exhibited essentially non existing levels of afucosylated N-glycans in M75-specific IgG1 Fc, which has been associated with exacerbated inflammatory responses and disease severity following viral infections ^44,45^. Our data provide evidence that sub-clinical exposure to *S. pyogenes* can selectively shape the N-glycan landscape of M75-specific IgG1 antibodies. The glycosylation features of the Fc region significantly influence the ability of IgG antibodies to trigger inflammatory responses and their interaction with FcγIII receptors ^46,47^. The concurrent increase in both agalactosylation and sialylation suggests a fine-tuned immune adaptation of M75-specific responses: the host may be enhancing IgG1 functionality for bacterial clearance (via agalactosylation ^48^) while simultaneously limiting collateral inflammation and tissue damage (via sialylation and fucosylation ^43,49^). This dual modulation could reflect an evolved mechanism to balance pathogen elimination with immune regulation during asymptomatic *S. pyogenes* exposures. Although the anti-M75 IgG response was predominantly mediated by the IgG1 and IgG3 subclasses post-challenge (as per ELISA data), the IgG Fc N-glycosylation in the non-pharyngitis (asymptomatic) participants appeared to be more prominent on IgG1 and IgG4 as compared to the pharyngitis group (both global and M75-IgG). Nevertheless, significant glycosylation changes were only observed in M75-specific IgG1 and not IgG3/4.

The modest immune responses to the conserved C-terminal M-protein peptide p*17 observed in this study are consistent with previous findings in both mice and children from streptococcal-endemic regions ^16^. Although C-terminal antigens appear to be cryptic during exposure and infection with *S. pyogenes*, vaccination with the p*17 antigen conjugated to a carrier protein elicits robust immune responses in animal models. These include high-affinity, class-switched antibodies that recognize a broad range of *S. pyogenes* strains, effectively reducing bacterial colonization and preventing severe infection ^23–26^. These observations underscore the likelihood that vaccine-induced protection against *S. pyogenes* operates through mechanisms distinct from naturally acquired immunity, which is only partially protective. This is evidenced by continued low susceptibility to infection among adults, particularly in parents of young children, individuals in crowded settings (e.g. army barracks), people who inject drugs, those experiencing homelessness, and older adults with accumulating comorbidities.

A limitation of our study is the small cohort size, especially the limited number of asymptomatic participants. However, these participants showed significant and strain-specific IgG binding to *S. pyogenes* equivalent to antibodies from mice immunised with M75 antigen. The high rate of clinical pharyngitis in the challenged adult participants, despite accumulated immunity to antigens other than the type-specific M75 hypervariable region (an exclusion criteria), may suggest the inoculum (dose) and direct swab challenge method may have circumvented or overwhelmed natural defences ^21^. Still, the inoculum was several orders of magnitude lower than in comparable small and larger animal models, around a third of challenged participants remained asymptomatic, and those participants did develop de novo type-specific antibodies. Beyond these concerns related to the model, this study did not consider responses to conserved non-M-protein antigens such as SpyCEP, SLO and ScpA, which have previously been found in this cohort to have been increased in participants with pharyngitis and to have persisted for at least 3 months post-challenge ^50^.

In conclusion, our study demonstrated that asymptomatic *S. pyogenes* infection can induce persistent serotype-specific functional antibodies. This finding could be re-examined in longitudinal *S. pyogenes* surveillance studies including serial serum sampling and tested in future human studies incorporating challenge and homologous rechallenge. We propose that sub-clinical exposures do contribute to the development of protective immunity against *S. pyogenes* in humans.

## Material and methods

### The CHIVAS-M75 study

The CHIVAS-M75 study protocol, including information on the M75 *S. pyogenes* challenge strain and key findings, has been detailed in earlier publications ^20–22^. Ethical approval for the study, including sample collection and associated immunological investigations, was granted by the Alfred Hospital Human Research Ethics Committee (reference number 500/17), and all participants provided written informed consent. Among the 25 participants enrolled in the trial, 19 developed pharyngitis, across two dose levels. The diagnosis of pharyngitis was corroborated by microbiological evidence (via qPCR and culture), elevated biochemical markers (C-reactive protein, cytokines, and chemokines), and serological responses (anti-streptolysin O and anti-DNase B antibodies). Importantly, all participants classified as asymptomatic in the present study showed no signs of *S. pyogenes* colonization. In the present study, analysis was limited by sample availability. Serum and PBMC were collected from participants at the evening prior to the challenge, then at 1-month and 6-month outpatient visits. Blood was collected in serum separator tubes (BD Vacutainer SST Gold 8.5 mL) and allowed to clot. Within 2 h of collection, tubes were centrifuged at 1500 × g for 15 min at 20 °C, and then aliquots were stored at −80 °C. PBMCs were collected using CPT tubes (BD Bioscience).

### Murine model of *S. pyogenes* infection

All animal protocols were reviewed and approved by the Griffith University Animal Ethics Committee (GU-AEC) in accordance with the National Health and Medical Research Council (NHMRC) of Australia guidelines. Experimental protocols involving SCID (Severe Combined Immunodeficiency) mice were reviewed and approved by Office of the Gene Technology Regulator (OGTR). Mice (female, 4–6 weeks) were sourced from Ozgene, Western Australia. All mice were housed in a PC2-certified animal facility in individually ventilated cages (IVC) with a maximum of 5 mice/cage. For positive controls in the pre-opsonization assay, Sprague Dawley rats (aged 9.5 weeks at the start of dosing) were sourced from Charles River Laboratories, Inc. The animals received three intramuscular injections of the p*17-K4S2-CRM/Alum vaccine over a six-week period, followed by a two-week recovery phase. The adjuvant, aluminium hydroxide (Alhydrogel 2%, Alum), was obtained from Brenntag Biosector, Denmark. The relative humidity ranged between 45–65% and temperature at 20–24 °C. Mice and rats were exposed to a 12-h light–dark cycle.

### Antigens

Synthetic peptides from the M75 protein hypervariable region (M75-HVR) and C-terminal region (p*17) regions were synthesized and purified (>95%) commercially by GenScript and stored lyophilized or in solution at −20 °C. Peptide sequence for M75-HVR is: EEERTFTELPYEARYKAWKSENDELRENYRRTLDKFNTEQGKTTRLEEQN. Peptide sequence for p*17 is: LRRDLDASREAKNQVERALE.

### Enzyme-linked immunosorbent assays (ELISA)

Standard ELISA was used to measure M75 and p*17-specific serum antibody levels. Briefly, peptides were coated onto NUNC MaxiSorp plates (Thermo Fisher). Antigens were resuspended to 5 µg/mL in 0.1 M coating carbonate buffer and coated at 100 µL per well onto 96-well medium-binding ELISA plates (Greiner Bio-One) overnight at room temperature. Plates were then blocked with 200 µL of 2% w/v bovine serum albumin (BSA, Sigma) in phosphate-buffered saline (PBS) overnight at 4 °C, followed by five washes in wash buffer (PBS with 0.05% v/v Tween20, PBS-T). Serial dilutions were performed using diluent (0.05% v/v Tween20) at 1:10, 1:20, 1:40, 1:80 dilutions. Samples were diluted 2-fold and antibody levels detected with HRP-conjugated goat anti-human IgA (Invitrogen, USA), IgG (H+L), IgG1, IgG2, IgG3 and IgG4 antibodies (Thermo Fisher, USA). SIGMAFAST OPD substrate was added according to manufacturer’s instructions and absorbance was measured at 450 nm on a Tecan Infinite M200 Pro plate reader (Tecan Group Ltd., Switzerland). Negative controls were included in each plate containing commercial human sera (Sigma) pre-absorbed of anti-M75 *S. pyogenes* antibodies or containing 0.05% v/v Tween20 only.

### Tetramer staining for peptide-specific memory B cells

Frozen mononuclear PBMCs were thawed at 37°C in thawing media (RPMI + 10% FCS), then washed and resuspended in flow buffer (PBS + 2%FCS + 2 mM EDTA). Biotinylated peptides synthesised at Genscript were tetramerised by serial addition of premium-grade streptavidin-PE and streptavidin-APC (Bio legend, San Diego, CA, USA) at a four-molar equivalent of biotinylated peptide to one molar equivalent of streptavidin during 2 h at room temperature ^51^. The prepared tetramer was diluted to a concentration of 2 μm for use. Typically, 10^6^ PBMCs were pre-incubated with Fc block (Bio legend) for 10 min followed by tetramer staining for 40 min. Cells were washed with flow buffer and then stained with fluorescently labelled antibodies (including CD19, CD20, CD21, CD27, IgD, IgM, IgG and IgA) and viability dye. Cells were fixed with 1% PFA and analyzed on a LSRFortessa (BD Biosciences). These cells were gated as positive for both fluorochromes of an antigen tetramer pair to reduce the inclusion of non-specific binding cells, as previously described ^51^.

### *S. pyogenes* strains and growth conditions

*S. pyogenes* strain 611024 (emm75, GenBank accession CP033621), isolated in Melbourne from a child presenting with exudative pharyngitis, was used in this study ^21^. *S. pyogenes* strain M1 5448 (emm1) was kindly provided by Professor Mark Walker, from the University of Queensland, isolated in Australia from a case of necrotising fasciitis. For mouse infection experiments, M75 *S. pyogenes* was serially passaged through mouse spleens to enhance virulence and rendered streptomycin-resistant (200 μg/mL) to enable selective recovery from commensal flora in respiratory mucosa and tissues.

*S. pyogenes* was cultured at 37 °C for 16–18 hours. Single colonies were inoculated into Todd Hewitt Broth (THB; Oxoid, Australia) supplemented with 1% Bacto Neopeptone (Thermo Fisher, USA), yeast extract and streptomycin ^24^. To enumerate colony-forming units (CFU), cultures were plated on Columbia agar containing 5% defibrinated horse blood and 200 μg/ml streptomycin (CBA 5%) ^24^.

### IgG purification

IgG antibodies were purified from the serum of CHIVAS participants and immunized rats using NAb Protein G columns, following the manufacturer’s instructions (Thermo Fisher, USA). The concentration of purified IgG was measured using a NanoDrop spectrophotometer (Thermo Fisher, USA).

M75 antigen-specific IgG antibodies were purified using SulfoLink™ affinity chromatography columns (Thermo Fisher Scientific, USA) following the manufacturer’s protocol. Briefly, the columns were prepared by coupling M75-HVR to the SulfoLink resin via sulfhydryl-reactive chemistry. Serum samples were then applied to the columns, allowing M75-HVR-specific IgG to bind selectively. After thorough washing to remove unbound proteins, antigen-specific IgG was eluted using a low pH glycine buffer and immediately neutralized. The purified IgG fractions were concentrated and buffer-exchanged into PBS using centrifugal filters (Amicon Ultra, Millipore). Purity of the eluted M75-specific IgG were assessed by silver protein stain (Thermo Fisher Scientific, USA) and ELISA. Purified antibodies were stored at –20°C until further use.

### IgG binding to *S. pyogenes*

For the binding assay, M1 and M75 *S. pyogenes* strains (0.5 O.D.) were incubated overnight in PBS containing 0.2% skim milk PBS with 100 µg of purified serum IgG collected from CHIVAS participants at pre-challenge and 6 months post-challenge. Following incubation, bacterial cells were washed and stained with anti-human IgG Fab antibody. Cells were fixed with 1% PFA and analyzed on a LSRFortessa (BD Biosciences) and FlowJo™ v11 Software was used for analysis.

### Pre-opsonization of *S. pyogenes* with IgG and intranasal challenge in mice

M75 *S. pyogenes* was incubated with purified serum IgG collected from asymptomatic CHIVAS participants at pre-challenge and 6 months post-challenge. As controls, IgG was purified from Sprague Dawley rats either immunised with p*17-DT-K4S2-CRM/Alum or left unimmunised (naïve). Following incubation, the bacterial cells were pre-incubated with a 10 μL inoculum containing 5 × 10^7^ CFU/mouse delivered intranasally to anaesthetized naive SCID mice. The protocol was a modification of a previous assay ^24^. Throat swabs were collected using flocked swabs (Copan Diagnostics, USA), which were then suspended in PBS, serially diluted, and plated in duplicate on CBA 5% plates. To assess nasal shedding, each mouse had its nares gently pressed onto a CBA 5% plates ten times (five times per half plate), allowing expelled particles to be streaked across the surface. On day 2 post-infection, mice were euthanized using CO₂ asphyxiation. The nasal-associated lymphoid tissue (NALT), a murine analogue to human tonsils, and the lungs were collected, homogenized in PBS using a Bullet Blender Homogenizer (Next Advance, USA), and serially diluted before being plated in duplicate onto CBA 5% plates. Mice were observed daily for clinical signs of illness according to a monitoring sheet approved by the GU-AEC.

### IgG Fc-glycopeptide profiling by LC-ESI MS/MS

Purified pre- and post-challenge global and M75 antigen-specific IgG from pharyngitis (n=5) and non-pharyngitis (asymptomatic) CHIVAS participants (n=5) were dried and redissolved in 25 mM ammonium bicarbonate buffer prior to trypsin digestion (enzyme-to-substrate ratio of 1:25) at 37°C. The resulting glycopeptides were analysed using an Orbitrap Fusion Tribrid mass spectrometer (Thermo Scientific), coupled to an UltiMate 3000 UHPLC fitted with a trap column (PepMap Neo C18, 5 mm x 300 µm, 5 µm particle size) and analytical reversed-phase C18 PepMap Neo (75 μm × 500 mm, 2 µm particle size) UHPLC column (Thermo Scientific) and PicoTip nanospray interface (New Objective). The samples were injected onto the trap column in 100 % Loading Buffer (0.1% trichloroacetic acid) at the flow rate of 15 µL/min and the (glyco)peptides were separated at a flow rate of 300 nL/min using linear gradient from 2% solvent A (0.1% formic acid) to 95% solvent B (80 % acetonitrile containing 0.1% formic acid) as follows: 1% solvent B until 8 min, 1%-8% from 8 to 9 min, 8-15 % from 9 to 20 min, 15-95% from 20 to 23 min, 95% from 23 to 25 min, before the column was re-equilibrated in 1% solvent B from 28 until 40 min. For MS acquisition, a full MS1 scan was performed at 60,000 resolution between m/z 350-1650 (custom AGC target at 250%, 50 ms maximum injection time, 1 microscan). Precursor ions were isolated using an isolation window of 1.6 Da and MS2 fragmentation was performed using normalized stepped-HCD collision energy (15, 30, 45%) at 60,000 resolution. All quantitative data processing was performed using Skyline (version 24.1) based on a pre-defined list of IgG1, IgG2, and IgG3/4 glycopeptide masses corresponding to doubly [M+2H]2+ and triply [M+3H]3+ charged precursor masses (Supp. Table 1). MS1 peak-area-under-the curve quantification was performed and the summed areas of integrated peaks (including monoisotopic transitions) for each glycopeptide precursor mass were represented as a percentage (relative intensity) of the total IgG glycopeptides.

### Statistical analysis

Statistical analysis was performed with GraphPad Prism 9 software using a nonparametric, unpaired Mann–Whitney U test (one-tailed; 90% confidence interval) to compare test groups, unpaired t test with Welch’s correction to compare test groups. ARRIVE guidelines ^52^ were used to calculate sample size for *in vivo* experiments. A sample size of 8 was shown to provide a power of 0.8 (G*Power) and therefore used for animal experiments involving bacterial challenge.

For ELISA analysis, absorbance values at a 1:20 serum dilution were used to compare antigen-specific responses across the three time points. Statistical analysis was conducted using one-way analysis of variance (ANOVA) with Geisser-Greenhouse correction to account for repeated measures. The same approach was applied to evaluate changes in antigen-specific responses over time in both the ELISA and antigen tetramer assays. Correlation coefficients and correlation p values were determined using Spearman’s method (two-sided) with a linear regression line included to indicate linear trends.

### Data availability

The IgG glycopeptide MS-raw data files have been made available via GlycoPost ^53^. For the reviewing process, the data is just available with the following information:

### ID: GPST000603, PIN CODE: 7330

URL:https://aus01.safelinks.protection.outlook.com/?url=https%3A%2F%2Fglycopost.glycosmos.org%2Fpreview%2F69820049068637b110aa34&data=05%7C02%7Cm.pandey%40griffith.edu.au%7Cd351e7fd918e453ee9b308ddb865c5be%7C5a7cc8aba4dc4f9bbf6066714049ad62%7C0%7C0%7C638869469437346273%7CUnknown%7CTWFpbGZsb3d8eyJFbXB0eU1hcGkiOnRydWUsIlYiOiIwLjAuMDAwMCIsIlAiOiJXaW4zMiIsIkFOIjoiTWFpbCIsIldUIjoyfQ%3D%3D%7C0%7C%7C%7C&sdata=xx1rushsW7asvF4%2Bot%2FLNw21qpYBxZAJNmZf0c8Du8c%3D&reserved=0

## Acknowledgement

The author thanks the support of the Flow Cytometry and Australian Cancer Research Foundation (ACRF) International Centre for Cancer Glycomics Mass Spectrometry Facility at the Institute for Biomedicine and Glycomics, Griffith University. A.L and A.C. are supported by an NHMRC project grant (APP1160379) awarded to M.P. V.O is supported by Griffith University Postgraduate Fellowship. M.A. is supported by an NHMRC ideas grant (GNT2018947) awarded to D.K. M.F.G. is supported by an NHMRC Investigator Fellowship (L3, GNT1174091). D.B. is supported by Queensland Advance WRAP grant awarded to A.L. The work described in this article was funded by The Heart Foundation of Australia (101656). The CHIVAS-M75 study was funded by the Australian National Health and Medical Research Council (1099183). J.O. is supported Australian National Health and Medical Research Council Investigator Fellowships and by a Melbourne Children’s Campus Clinician–Scientist fellowship. H.R.F. and K.I.A. are Human Infection Challenge Network for Vaccine Development members.

## Author contributions

Conceptualization: A.L., M.F.G., M.P., D.K., A.C.S. and J.O. Methodology and experiments: A.L., D.V., V.O., M.A., D.B., A.C, H.R.F. and K.I.A. Data analysis: A.L., D.V., D.B. M.A., Patient cohorts: A.C.S. and J.O. Original draft: A.L., M.F.G. and M.P. Manuscript review: A.L., D.V., V.O., K.I.A., J.O., M.P., D.K. and M.F.G. Funding acquisition: A.C.S., J.O., A.L., D.K., M.P. and M.F.G.

