## Supplementary material for "Sub-clinical exposure to *Streptococcus pyogenes* drives the development of immunity": Supp figures and Table

### Supplementary Figure 1

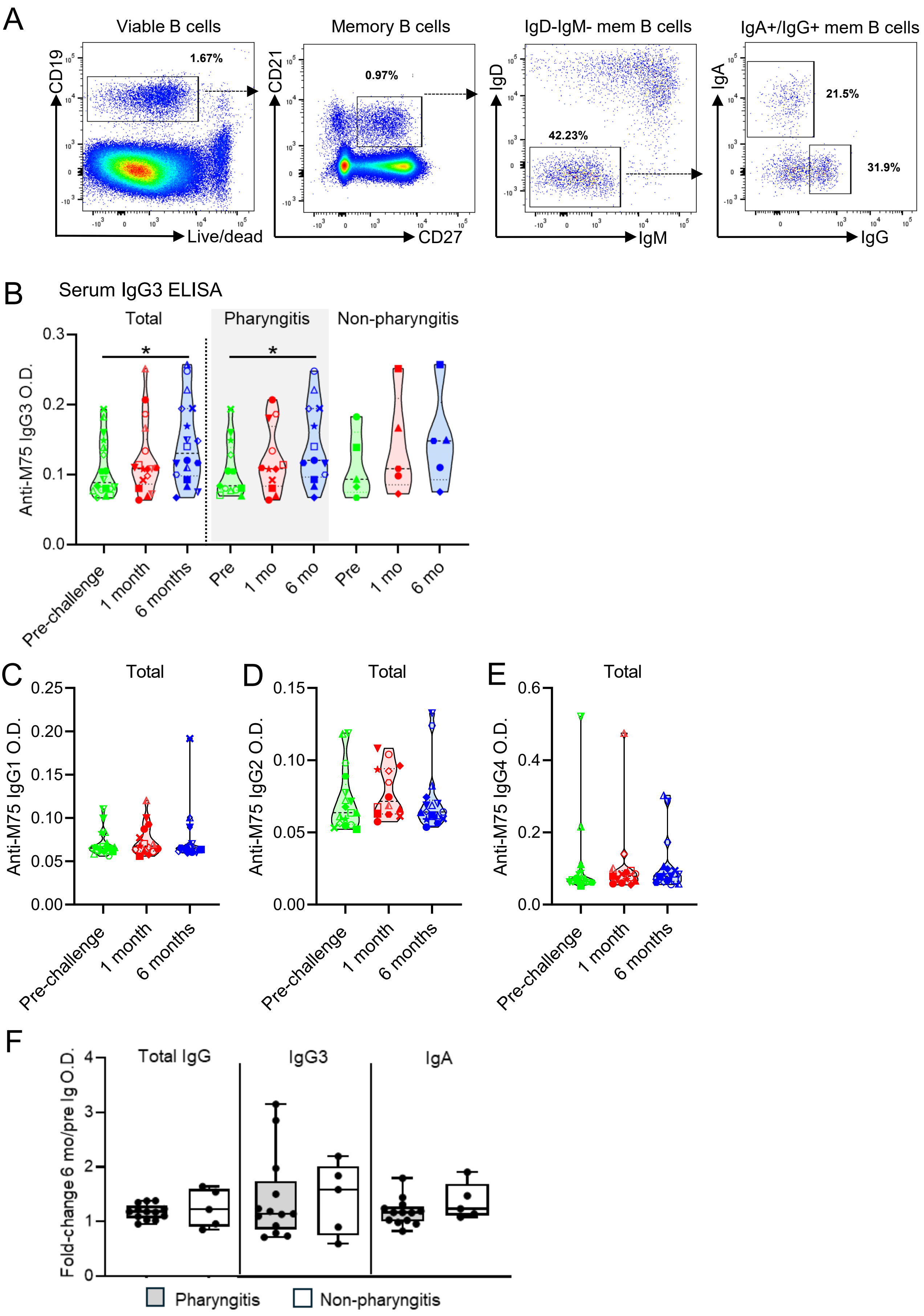

**Supplementary Figure 1.** Antigen-specific immune responses to M75-HVR following experimental human Strep A M75 infection. (A) Flow cytometry gating strategy for analysis of $IgG^+$ and $IgA^+$ memory B cells in the peripheral blood. (B,C) ) ELISA was used to detect anti-M75-specific IgG antibody subtypes in serum samples. Violin plots show O.D. 450 absorbance values are shown for sera collected at pre-challenge (day -1), 1 month post-challenge and 6 months post-challenge, using a 1:20 dilution. Violine plots show the absorbance values of M75-specific (IgG3 (C) IgG1, (D) IgG2 and (E) IgG4. Violin plots show O.D. 450 absorbance values for sera collected at pre-challenge (day -1) and 6 months post-challenge, using a 1:20 dilution. Data are stratified by participants who developed pharyngitis and those who did not. Each individual is represented by a symbol. One-way ANOVA followed by Bonferroni's comparisons tests were performed in all statistical analyses. *$P<0.05$, **$P<0.01$. (F) Box and whiskers plot of the ELISA data showing OD fold-change (6 months/ pre-infection) for M75-specific total IgG, IgG3, and IgA at a 1:20 dilution. Median and 25th, and 75th percentiles (boxes), as well as the minimum and maximum values, are indicated. The fold change represents the difference in OD values of 6 months post-infection compared to pre-infection levels.

### Supplementary Figure 2

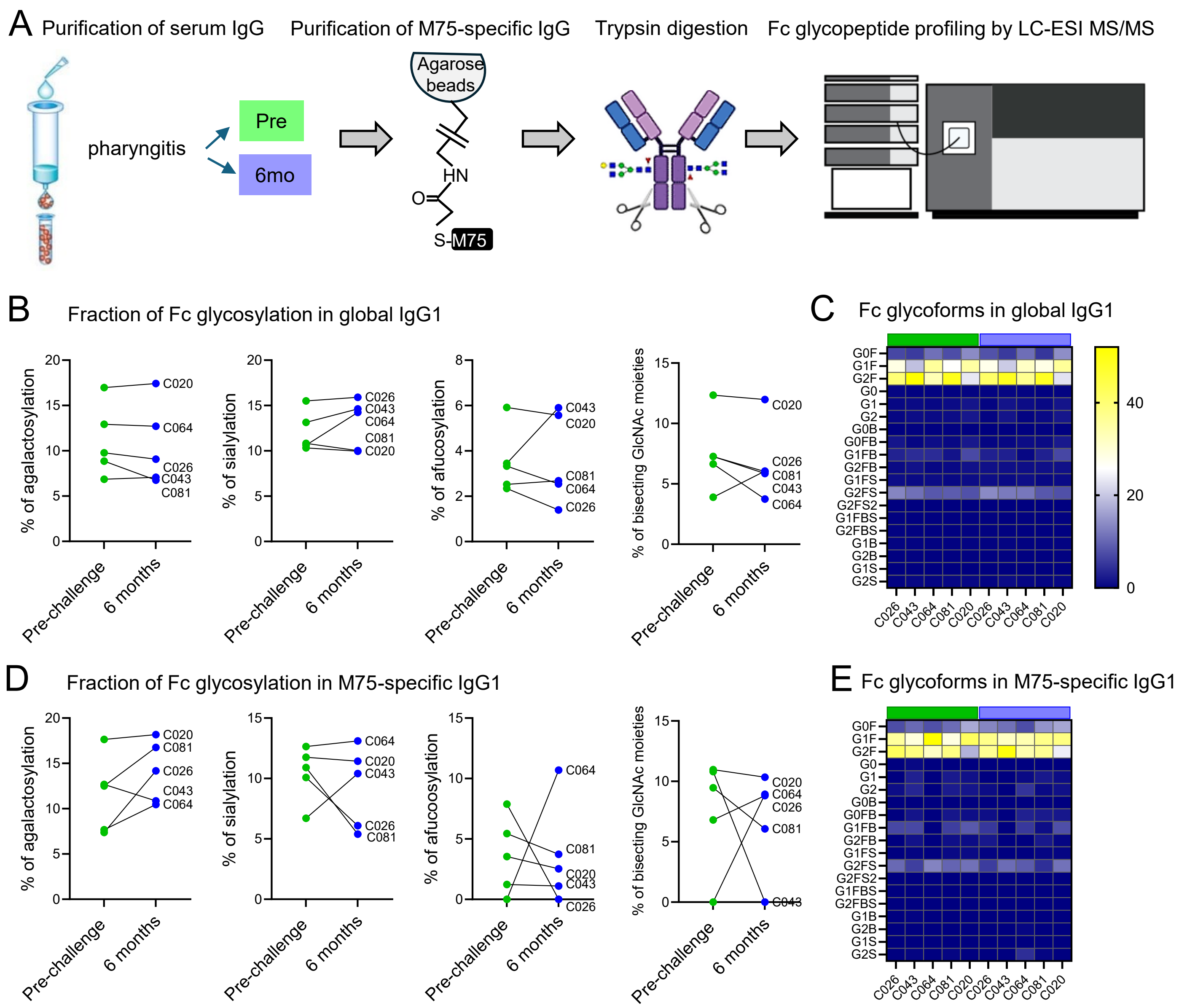

**Supplementary Figure 2**. Fc N-glycosylation profiling of global and M75-specific serum IgG1 pre- and 6 months post-challenge in participants with symptomatic pharyngitis. (A) Schematic overview of M75-specific IgG purification, digestion and Fc glycan profiling by LC-ESI MS/MS. (B) Line graphs depict the relative abundance of glycans in global IgG1 antibodies. (C) Heatmaps illustrate the fractional distribution of individual Fc glycoforms in global IgG1 antibodies. (D) Line graphs depict the relative abundance of glycans in M75-specific antibodies. (E) Heatmaps illustrate the fractional distribution of Fc glycoforms in M75-specific IgG1 antibodies. Data represent paired samples collected at the time of challenge (green) and 6 months post-challenge (blue) from participants with pharyngitis (n = 5). Paired t-test between pre- and post-challenge was used for statistical analysis. Statistical analysis revealed no significant differences between the groups.

| Retention Time | Glycan ID | Glycan Structure | Precursor Mass | |
|---|---|---|---|---|
| | | | [M+3H]$^{3+}$ | [M+2H]$^{2+}$ |
| **IgG1** | | | | |
| 23.5-24.5 | IgG1_G0F | Hex[3]HexNAc[4]Fuc[1] | 878.6868 | 1317.5266 |
| 23.5-24.5 | IgG1_G1F | Hex[4]HexNAc[4]Fuc[1] | 932.7044 | 1398.5530 |
| 23.5-24.5 | IgG1_G2F | Hex[5]HexNAc[4]Fuc[1] | 986.7220 | 1479.5794 |
| 23.5-24.5 | IgG1_G0 | Hex[3]HexNAc[4] | 830.0008 | 1244.4976 |
| 23.5-24.5 | IgG1_G1 | Hex[4]HexNAc[4] | 884.0184 | 1325.5240 |
| 23.5-24.5 | IgG1_G2 | Hex[5]HexNAc[4] | 938.0361 | 1406.5504 |
| 23.5-24.5 | IgG1_G0B | Hex[3]HexNAc[5] | 897.6940 | 1346.0373 |
| 23.5-24.5 | IgG1_G0FB | Hex[3]HexNAc[5]Fuc[1] | 946.3799 | 1419.0663 |
| 23.5-24.5 | IgG1_G1FB | Hex[4]HexNAc[5]Fuc[1] | 1000.3975 | 1500.0927 |
| 23.5-24.5 | IgG1_G2FB | Hex[5]HexNAc[5]Fuc[1] | 1054.4151 | 1581.1191 |
| 25.4-26.0 | IgG1_G1FS | Hex[4]HexNAc[4]Fuc[1]NeuAc | 1029.7362 | 1544.1007 |
| 25.4-26.0 | IgG1_G2FS | Hex[5]HexNAc[4]Fuc[1]NeuAc | 1083.7538 | 1625.1271 |
| 25.4-26.0 | IgG1_G2FS2 | Hex[5]HexNAc[4]Fuc[1]NeuAc[2] | 1180.7856 | 1770.6748 |
| 25.4-26.0 | IgG1_G1FBS | Hex[4]HexNAc[5]Fuc[1]NeuAc | 1097.4293 | 1645.6404 |
| 25.4-26.0 | IgG1_G2FBS | Hex[5]HexNAc[5]Fuc[1]NeuAc | 1151.4470 | 1726.6668 |
| 23.5-24.5 | IgG1_G1B | Hex[4]HexNAc[5] | 951.7116 | 1427.0637 |
| 23.5-24.5 | IgG1_G2B | Hex[5]HexNAc[5] | 1005.7292 | 1508.0901 |
| 25.4-26.0 | IgG1_G1S | Hex[4]HexNAc[4]NeuAc | 981.0503 | 1471.0717 |
| 25.4-26.0 | IgG1_G2S | Hex[5]HexNAc[4]NeuAc | 1035.0679 | 1552.0982 |
| **IgG2** | | | | |
| 28.0-29.5 | IgG2_G0F | Hex[3]HexNAc[4]Fuc[1] | 868.0235 | 1301.5317 |
| 28.0-29.5 | IgG2_G1F | Hex[4]HexNAc[4]Fuc[1] | 922.0411 | 1382.5581 |
| 28.0-29.5 | IgG2_G2F | Hex[5]HexNAc[4]Fuc[1] | 976.0587 | 1463.5845 |
| 28.0-29.5 | IgG2_G0 | Hex[3]HexNAc[4] | 819.3376 | 1228.5027 |
| 28.0-29.5 | IgG2_G1 | Hex[4]HexNAc[4] | 873.3552 | 1309.5291 |
| 28.0-29.5 | IgG2_G2 | Hex[5]HexNAc[4] | 927.3728 | 1390.5555 |
| 28.0-29.5 | IgG2_G0B | Hex[3]HexNAc[5] | 887.0307 | 1330.0424 |
| 28.0-29.5 | IgG2_G0FB | Hex[3]HexNAc[5]Fuc[1] | 935.7167 | 1403.0713 |
| 28.0-29.5 | IgG2_G1FB | Hex[4]HexNAc[5]Fuc[1] | 989.7343 | 1484.0978 |
| 28.0-29.5 | IgG2_G2FB | Hex[5]HexNAc[5]Fuc[1] | 1043.7519 | 1565.1242 |
| 29.5-31 | IgG2_G1FS | Hex[4]HexNAc[4]Fuc[1]NeuAc | 1019.0729 | 1528.1058 |
| 29.5-31 | IgG2_G2FS | Hex[5]HexNAc[4]Fuc[1]NeuAc | 1073.0906 | 1609.1322 |
| 29.5-31 | IgG2_G2FS2 | Hex[5]HexNAc[4]Fuc[1]NeuAc[2] | 1170.1224 | 1754.6799 |
| 29.5-31 | IgG2_G1FBS | Hex[4]HexNAc[5]Fuc[1]NeuAc | 1086.7661 | 1629.6455 |
| 29.5-31 | IgG2_G2FBS | Hex[5]HexNAc[5]Fuc[1]NeuAc | 1140.7837 | 1710.6719 |
| 28.0-29.5 | IgG2_G1B | Hex[4]HexNAc[5] | 941.0483 | 1411.0688 |
| 28.0-29.5 | IgG2_G2B | Hex[5]HexNAc[5] | 995.0659 | 1492.0952 |
| 29.5-31 | IgG2_G1S | Hex[4]HexNAc[4]NeuAc | 970.3781 | 1455.0768 |
| 29.5-31 | IgG2_G2S | Hex[5]HexNAc[4]NeuAc | 1024.4046 | 1536.1032 |
| **IgG3/4** | | | | |
| 26.2-27 | IgG3/4_G0F | Hex[3]HexNAc[4]Fuc[1] | 873.3552 | 1309.5291 |
| 26.2-27 | IgG3/4_G1F | Hex[4]HexNAc[4]Fuc[1] | 927.3728 | 1390.5555 |
| 26.2-27 | IgG3/4_G2F | Hex[5]HexNAc[4]Fuc[1] | 981.3904 | 1471.5819 |
| 26.2-27 | IgG3/4_G0 | Hex[3]HexNAc[4] | 824.6692 | 1236.5002 |
| 26.2-27 | IgG3/4_G1 | Hex[4]HexNAc[4] | 878.6868 | 1317.5266 |
| 26.2-27 | IgG3/4_G2 | Hex[5]HexNAc[4] | 932.7044 | 1398.553 |
| 26.2-27 | IgG3/4_G0B | Hex[3]HexNAc[5] | 892.3623 | 1338.0399 |
| 26.2-27 | IgG3/4_G0FB | Hex[3]HexNAc[5]Fuc[1] | 941.0483 | 1411.0688 |

| | | | | |
|---|---|---|---|---|
| 26.2-27 | IgG3/4_G1FB | Hex[4]HexNAc[5]Fuc[1] | 995.0659 | 1492.0952 |
| 26.2-27 | IgG3/4_G2FB | Hex[5]HexNAc[5]Fuc[1] | 1049.0835 | 1573.1216 |
| 29-30 | IgG3/4_G1FS | Hex[4]HexNAc[4]Fuc[1]NeuAc | 1024.4046 | 1536.1032 |
| 29-30 | IgG3/4_G2FS | Hex[5]HexNAc[4]Fuc[1]NeuAc | 1078.4222 | 1617.1297 |
| 29-30 | IgG3/4_G2FS2 | Hex[5]HexNAc[4]Fuc[1]NeuAc[2] | 1175.454 | 1762.6774 |
| 29-30 | IgG3/4_G1FBS | Hex[4]HexNAc[5]Fuc[1]NeuAc | 1092.0977 | 1637.6429 |
| 29-30 | IgG3/4_G2FBS | Hex[5]HexNAc[5]Fuc[1]NeuAc | 1146.1153 | 1718.6693 |
| 26.2-27 | IgG3/4_G1B | Hex[4]HexNAc[5] | 946.3799 | 1419.0663 |
| 26.2-27 | IgG3/4_G2B | Hex[5]HexNAc[5] | 1000.3975 | 1500.0927 |
| 29-30 | IgG3/4_G1S | Hex[4]HexNAc[4]NeuAc | 975.7186 | 1463.0743 |
| 29-30 | IgG3/4_G2S | Hex[5]HexNAc[4]NeuAc | 1029.7362 | 1544.1007 |

Table 1: List of IgG precursor MS1 glycopeptide masses depicted as doubly $[M+2H]^{2+}$ and triply $[M+3H]^{3+}$ charged masses used for Skyline quantification. Retention times used for MS1 precursor selection and integration of area-under-the curve are indicative of retention time shifts observed using C18 column. IgG samples from all non-pharyngitis (n=5) and pharyngitis (n=4) patients were analysed over a 40 min LC-MS gradient; except for one pharyngitis sample (72 min LC-MS gradient). MS1 precursors selected for quantification were manually verified based on presence of glycopeptide-specific backbone ion [Pep+HexNAc: Y1_IgG1: *m/z* $1392.5914^{1+}$; Y1_IgG2: *m/z* $1360.6016^{1+}$ and Y1_IgG3/4: *m/z* $1376.5965^{1+}$) and oxonium ion (*m/z* $204.0869^{1+}$).
